# Structural Mapping of Polyclonal IgG Responses to HA After Influenza Virus Vaccination or Infection

**DOI:** 10.1101/2024.07.08.601940

**Authors:** André Nicolás León, Alesandra J. Rodriguez, Sara T. Richey, Alba Torrents de la Peña, Rachael M. Wolters, Abigail M. Jackson, Katherine Webb, C. Buddy Creech, Sandra Yoder, Philip A. Mudd, James E. Crowe, Julianna Han, Andrew B. Ward

**Author notes:** Correspondence should be addressed to J.H. and A.B.W.

## Abstract

Cellular and molecular characterization of immune responses elicited by influenza virus infection and seasonal vaccination have informed efforts to improve vaccine efficacy, breadth, and longevity. Here, we use negative stain electron microscopy polyclonal epitope mapping (nsEMPEM) to structurally characterize the humoral IgG antibody responses to hemagglutinin (HA) from human patients vaccinated with a seasonal quadrivalent flu vaccine or infected with influenza A viruses. Our data show that both vaccinated and infected patients had humoral IgGs targeting highly conserved regions on both H1 and H3 subtype HAs, including the stem and anchor, which are targets for universal influenza vaccine design. Responses against H1 predominantly targeted the central stem epitope in infected patients and vaccinated donors, whereas head epitopes were more prominently targeted on H3. Responses against H3 were less abundant, but a greater diversity of H3 epitopes were targeted relative to H1. While our analysis is limited by sample size, on average, vaccinated donors responded to a greater diversity of epitopes on both H1 and H3 than infected patients. These data establish a baseline for assessing polyclonal antibody responses in vaccination and infection, providing context for future vaccine trials and emphasizing the importance of carefully designing vaccines to boost protective responses towards conserved epitopes.

## Introduction

The 1918 H1N1 flu pandemic killed more people than armed conflict in WWI, reshaping labor markets and catalyzing fundamental changes in U.S. public health policy (1, 2). Over 90 years later, individuals who survived H1N1 infection in the early 20^th^ century were found to express antibodies that improved their ability to combat infection by the antigenically-similar H1N1 swine flu that began circulating in 2009 (3, 4). Given its clear relevance to influenza pathogenesis, vaccine design, and basic immunology, the molecular basis and manifestation of this long-lived immune memory has been a topic of intense study (5–8).

Humoral immunity to influenza viruses primarily targets two viral surface glycoproteins: hemagglutinin (HA) and neuraminidase (NA) (9). HA is a homotrimeric, archetypical Type-I membrane fusion protein that binds to sialic acid containing glycans on the surface of host cells with differential binding affinities for avian and human sialic acid linkages (10). NA is a potent, homotetrameric sialidase that facilitates escape from highly sialylated mucins in the mucosa and allows budding viruses to release from the host cell surface, thereby preventing reentry into previously infected cells (11). Influenza A virus (IAV) HA comes in 19 different subtypes (H1-H19), NA in 11 (N1-N11). Subtypes are distinguished by sequence identity and vary in their antigenicity, activity, and receptor specificities. Influenza viruses are thus classified by their HA and NA subtype pairings. H1N1 and H3N2 are currently the clinically relevant IAV subtypes circulating in humans.

Given its vital roles in receptor binding and membrane fusion, high concentration on the viral surface, relative ease of production, and immunodominance, HA has been the principal target of vaccine design efforts (5, 12–15). As such, the hemagglutinin inhibition assay (HAI) has been the gold-standard correlate of protection for evaluating vaccine efficacy, and only HA concentrations are controlled in seasonal vaccine formulations. However, due to immune pressure, HA undergoes extensive mutagenesis and is frequently subject to antigenic drift due to the accumulation of mutations in the immunodominant head region. This antigenic drift necessitates global surveillance of influenza sequences to predict what strains to include in yearly vaccines. Even with modern surveillance infrastructure, mismatches between circulating and vaccine strains can and do occur, which can cause vaccine efficacies to decrease precipitously (16–18). Recombination events between different influenza virus subtypes can also lead to antigenic shifts where new HAs begin to circulate in humans. The pandemic potential of recombination events demands the development of vaccination programs that far exceed the efficacy of current seasonal vaccine cocktails with the goal of developing a universal influenza vaccine.

Compared to the immunodominant head, the HA central stem and anchor epitopes are relatively conserved, and recent works characterizing broadly neutralizing antibodies targeting these regions highlight their viability as targets for a universal influenza vaccine (19–21). In contrast to head antibodies that inhibit receptor engagement, protective central stem and anchor antibodies can function by blocking membrane fusion, eliciting Fc-mediated effector function, and engaging with serum complement protein C1q (9, 22). Creative antigen engineering efforts to bias responses to conserved epitopes in the head and central stem have been tested in animal models and human clinical trials with promising results (20, 21, 23–27). Much of this work has benefitted from the extensive structural characterization of antibodies bound to HA.

Since the first structures of HA were reported by Wilson and Wiley in 1983, hundreds of structures of HA and HA-antibody complexes have been deposited to the PDB (28–30). As crystallography and cryo-electron microscopy techniques became more accessible, the pace of deposition rapidly increased, providing a wealth of structural information for vaccine design (31, 32). While high-resolution structures are invaluable, low-resolution data can also prove informative, particularly for mapping highly diverse polyclonal antibody responses. Electron microscopy polyclonal epitope mapping (EMPEM) uses low-resolution negative stain EM images to map epitopes targeted by heterogeneous polyclonal antibodies purified from sera (33). EMPEM allows the mapping of multiple epitopes in a single mixture, unlike competitive ELISAs, which require control panels of well-characterized monoclonal antibodies which can introduce false-positives/negatives due to steric hindrance and out-competing low-affinity antibodies present in sera. Recent reports have used EMPEM to map epitopes targeted by human patients vaccinated against or infected with various infectious diseases, including HIV-1, SARS-CoV-2, and IAV (34–37). EMPEM has also been used to identify responses to novel epitopes, which has already led to the characterization of a new broadly neutralizing HA epitope, the aforementioned anchor epitope (19).

In this study, EMPEM was used to map serum antibody responses against both H1 and H3 HAs from human donors who had been vaccinated with a quadrivalent vaccine or naturally infected with an IAV. This work aims to establish a baseline for influenza EMPEM to facilitate continued use in investigations into induced and recalled responses during infection and vaccination and as an assay in large-scale vaccination trials. To that end, our study found that 1) antigen^+^ responses are highly heterogeneous across individuals and do not neatly sort into subgroups, and 2) responses to the highly conserved central stem were almost ubiquitous in both infected patients and vaccinated donors.

## Results

### Infected patients demonstrated ubiquitous responses to H1 and H3 central stem

Patients (n = 7) presented to the emergency department reporting symptoms consistent with an acute respiratory virus infection. Subsequent testing confirmed infection with H1N1 (n = 4) or H3N2 (n = 3) IAV viruses. Patient ages ranged from 18 to 73 years old. As patients were not infected in a controlled influenza virus challenge, it is not possible to know exactly when they were infected; however, self-reported days symptomatic were recorded and suggested that patients presented within the first week of infection (**Table 1**).

**Table 1:**
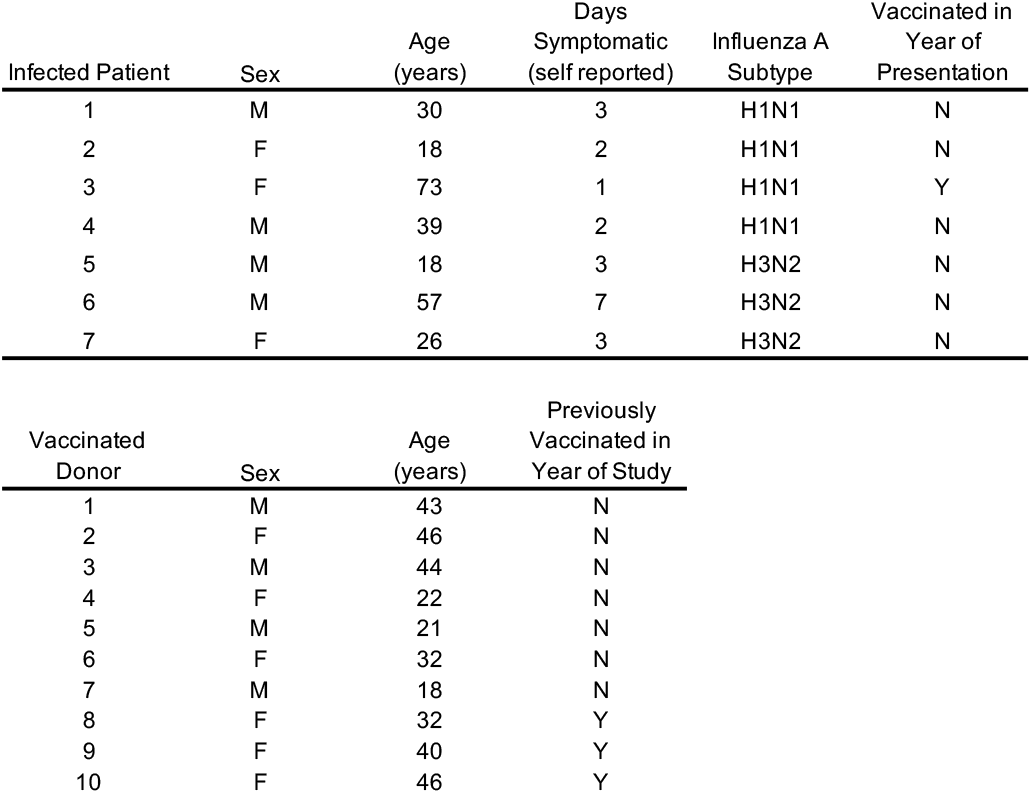
Characteristics of study participants by method of antigen exposure during 2018-2019 influenza season.

To assess the HA epitopes (**Figure 1)** targeted by antibodies at the time of presentation, IgG was purified from the sera of infected patients, digested into Fab, and complexed with either A/Michigan/45/2015 (H1) or A/Singapore/INFIMH-009/2016 (H3) (**Figure 2, S1**) (10). EMPEM revealed that all patients had central stem responses to both H1 and H3. Patients 1, 2, 3, 5, and 7 had responses to the H1 head, and Patients 1, 2, 3, 4, and 7 had responses to the H3 head. Only Patients 2, 3, and 5 showed responses to the side of the H3 head or vestigial esterase epitope.

**Figure 1:**
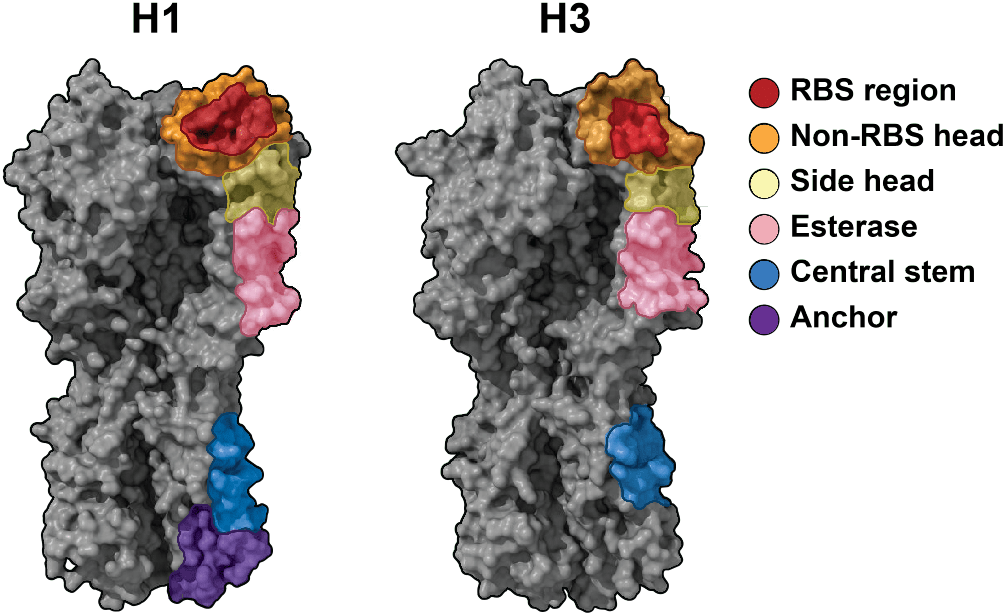
Epitope map of HA responses. Representative volumes for H1 (PDB: 4M4Y) and H3 (PDB: 4ZCJ) with epitope footprints represented (10).

**Figure 2:**
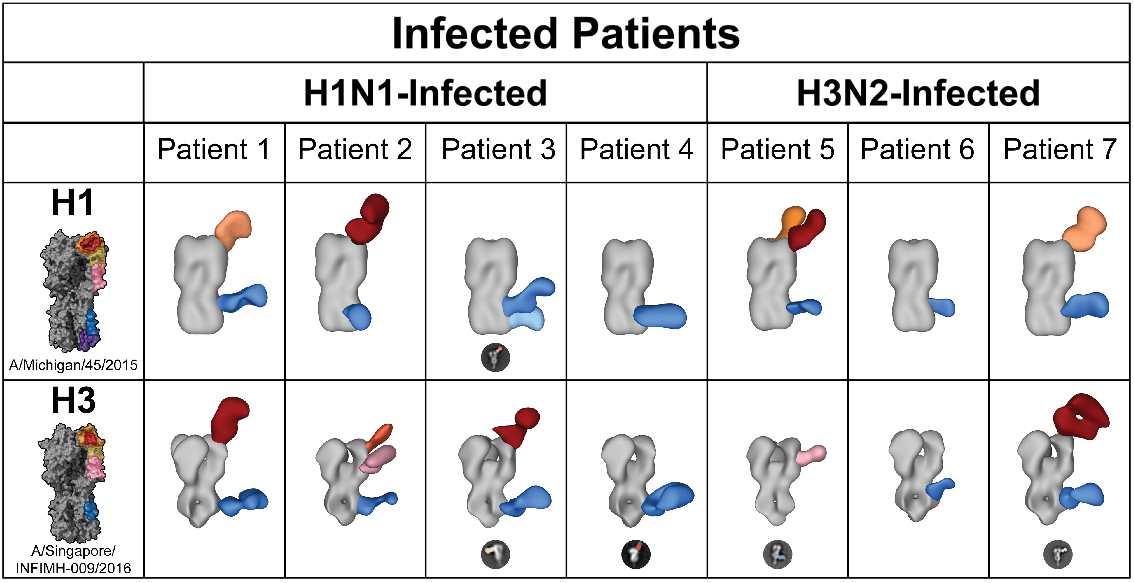
HA^+^ polyclonal IgG Fab responses present in infected patient sera at time of presentation. Negative stain EM reconstructions of purified polyclonal IgG Fab bound to A/Michigan/45/2015 (H1) or A/Singapore/INFIMH-009/2016 (H3). Representative, false-colored 2D classes are presented for epitopes that could not be 3D reconstructed.

These data indicate that at the time of presentation, all but two patients (4 and 6) had detectable humoral IgG targeting multiple HA epitopes and that all patients targeted the conserved central stem. To contextualize these results and establish a better understanding of epitope dynamics over time, EMPEM was conducted on sera of vaccinated donors in a controlled study.

### Vaccinated donors had a greater diversity of responses to H1 and H3

Ten donors were enrolled in a study to assess the dynamics of epitope targeting over time after vaccination with a quadrivalent flu vaccine (**Table 1**). Donors were vaccinated with the 2018-2019 Flucelvax Quadrivalent formulation containing A/Singapore/GP1908/2015 IVR-180 (H1), A/North Carolina/04/2016 (H3), and representative strains for influenza B/Yamagata and B/Victoria lineages. Sera were collected on Day 0 and Day 14 and processed for EMPEM analysis (**Figure 3, S2-3**). As the vaccine was administered at a known timepoint, it is feasible to make more substantiated hypotheses about the immunological origin of the observed responses.

**Figure 3:**
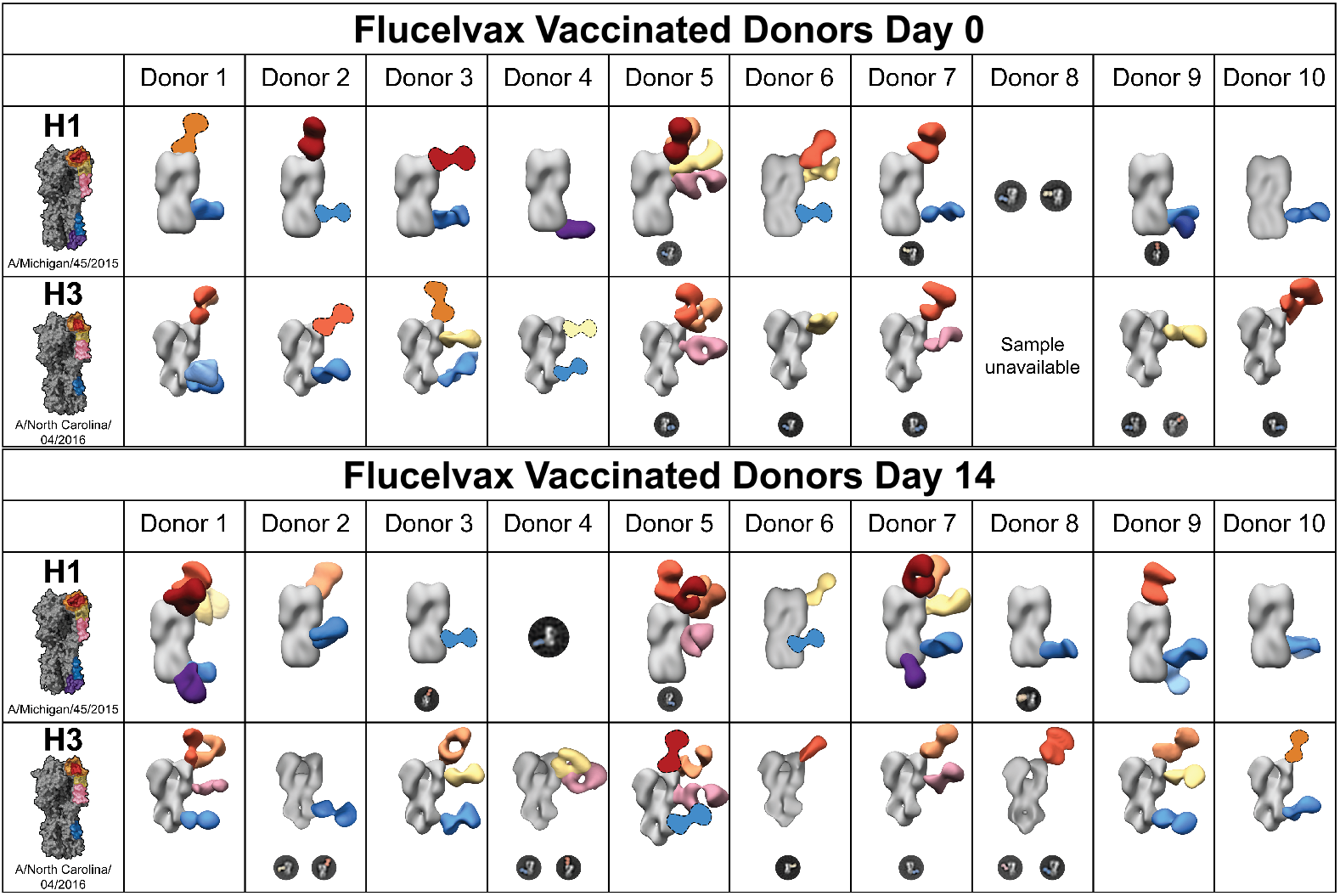
HA^+^ polyclonal IgG Fab responses before and after vaccination with quadrivalent Flucelvax. Negative stain EM reconstructions of purified polyclonal IgG Fab bound to A/Michigan/45/2015 (H1) or A/North Carolina/04/2016 (H3). Representative, false-colored 2D classes are presented for epitopes that could not be 3D reconstructed. 3D reconstructions that illustrated epitopes targeted but were poorly resolved are presented with cartoon Fabs. Donors were vaccinated with the 2018-2019 FLUCELVAX QUADRIVALENT formulation containing A/Singapore/GP1908/2015 IVR-180 (H1), A/North Carolina/04/2016 (H3), and representative strains for influenza B/Yamagata and B/Victoria lineages.

Donors targeted anywhere from 1 to 5 distinct epitopes at Day 0; all donors had responses to the H1 and H3 central stem except Donor 4, who had a response to the anchor epitope of H1 but no observable central stem response. Central stem responses were also nearly ubiquitous on Day 14, except for Donor 6 to H3. Donor 5 had a particularly diverse response to both H1 and H3 on Day 0 and Day 14, presenting six distinct responses to H1 on Day 14, including four unique responses to the head. Donors 1 and 3 also targeted a relatively high diversity of epitopes on H3, including multiple central stem epitopes by Donor 1 at Day 0.

The main differentiating factor between vaccinated donors at Day 0 and the infected patients was not necessarily the number of epitopes targeted but which epitopes were targeted. Whereas the infected patients almost exclusively targeted the central stem, RBS, and non-RBS head, the vaccinated donors targeted the side head, vestigial esterase, and anchor with higher frequency.

Only Donors 1, 5, and 7 showed an increase in the number of epitopes targeted on Day 14 compared to Day 0. Donors 1 and 7 targeted at least five distinct epitopes on H1 on Day 14 compared to 2-3 on Day 0. Donor 5 exhibited a more modest increase from 5 H1 epitopes to 6. However, 7/10 donors targeted different H1 epitopes from Day 0 to Day 14, and 5/9 followed a similar trend with H3.

Given these observations on the number and identity of epitopes targeted at baseline, during the early response to infection, and at the peak of germinal center (GC) mediated responses, serological analyses were conducted to probe for functional correlations.

### Assessing trends between EMPEM and serology

Diluted plasma was used to conduct ELISAs against HAs from A/Michigan/45/2015 and A/Singapore/INFIMH-009/2016 and analyzed to calculate the area under the curve (AUC) (**Figure 4A-B**). These ELISAs demonstrated a 2.5- to 3-fold increase in AUC from Day 0 to Day 14 in vaccinated donors for H1 and H3, consistent with clonal expansion and proliferation of affinity matured, GC-derived plasma cells. AUCs for infected patients were comparable to vaccinated donors at Day 0 and 4-fold lower than vaccinated donors at Day 14.

**Figure 4:**
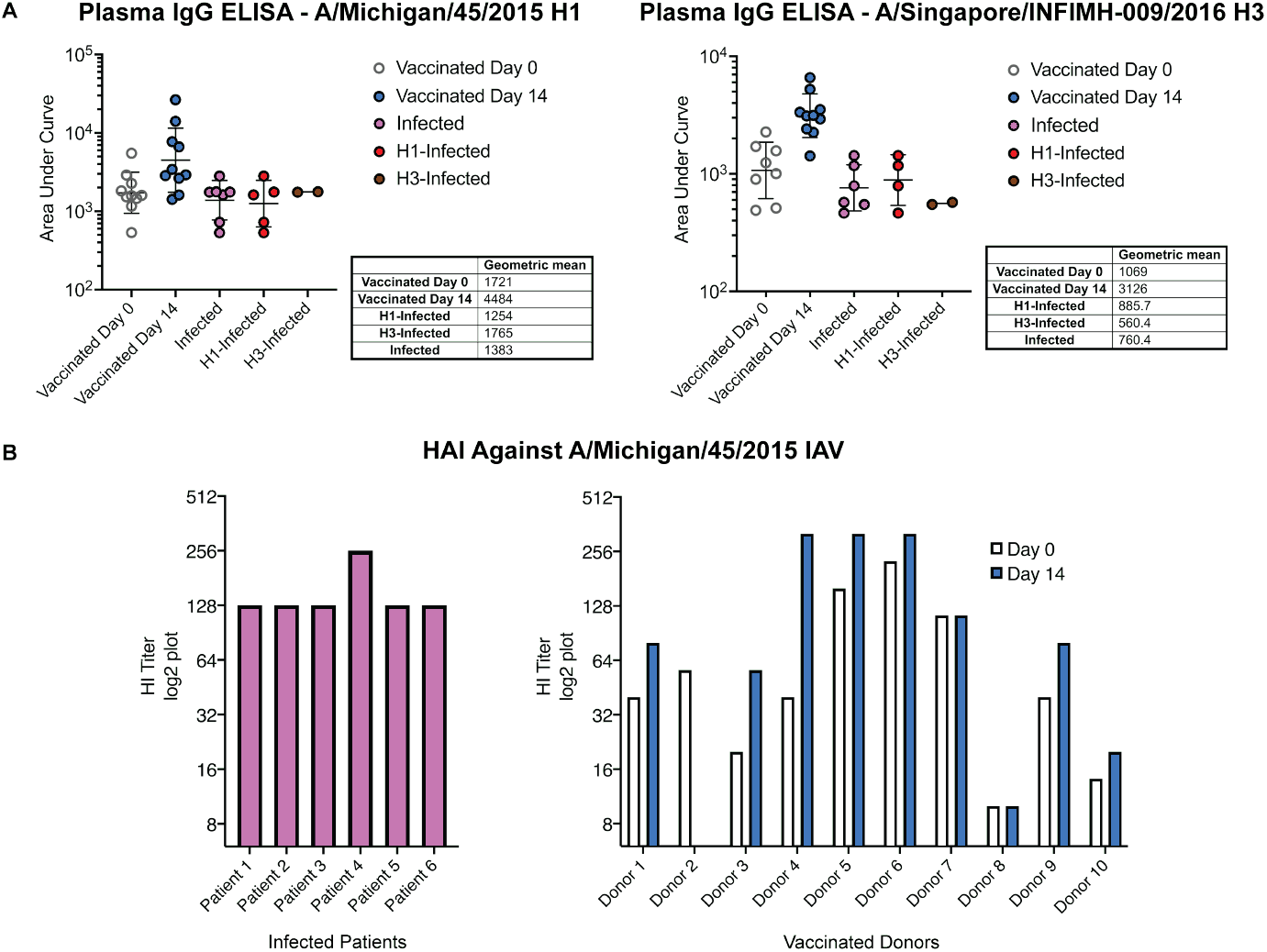
Greater number of HA^+^ polyclonal IgG responses trend with greater binding via ELISA but not increase HI titer. **(A)** HA reactivity across groups was measured by plasma ELISA and reported as AUC and geometric means. **(B)** HA inhibition assays were conducted using patient plasma and are presented as log2 plots.

Hemagglutinin inhibition (HI) assays were also conducted against A/Michigan/45/2015 virus propagated in eggs. Despite lacking observable responses targeting the H1 RBS or head, both Patient 4 and Patient 6 had HI titers comparable to the remaining patients, suggesting that the stem-targeting IgGs observed in EMPEM may inhibit hemagglutination through steric hindrance or that RBS-targeting antibodies were present in their sera but unable to be detected through EMPEM (**Figures 2, 4**). This finding could be due to the loss of avidity after IgG digestion to Fab or that non-IgG antibodies present in serum (*i*.*e*., IgA and IgM) are not accounted for in the standard EMPEM workflow. Complement component C1q may also contribute to HI observed in the absence of head- or RBS-targeting antibodies (22).

To explore this finding further, the particle distribution profiles of the nsEMPEM data sets were analyzed (**Figures 5-6:B-C**). Consistent with ELISA trends for H1 and H3, infected patients and vaccinated donors had a higher proportion of unbound H3 than H1 on average. Indeed, particle distributions matched well with all ELISA trends previously highlighted, but they had similar limitations in their ability to explicate trends observed in HI.

**Figure 5:**
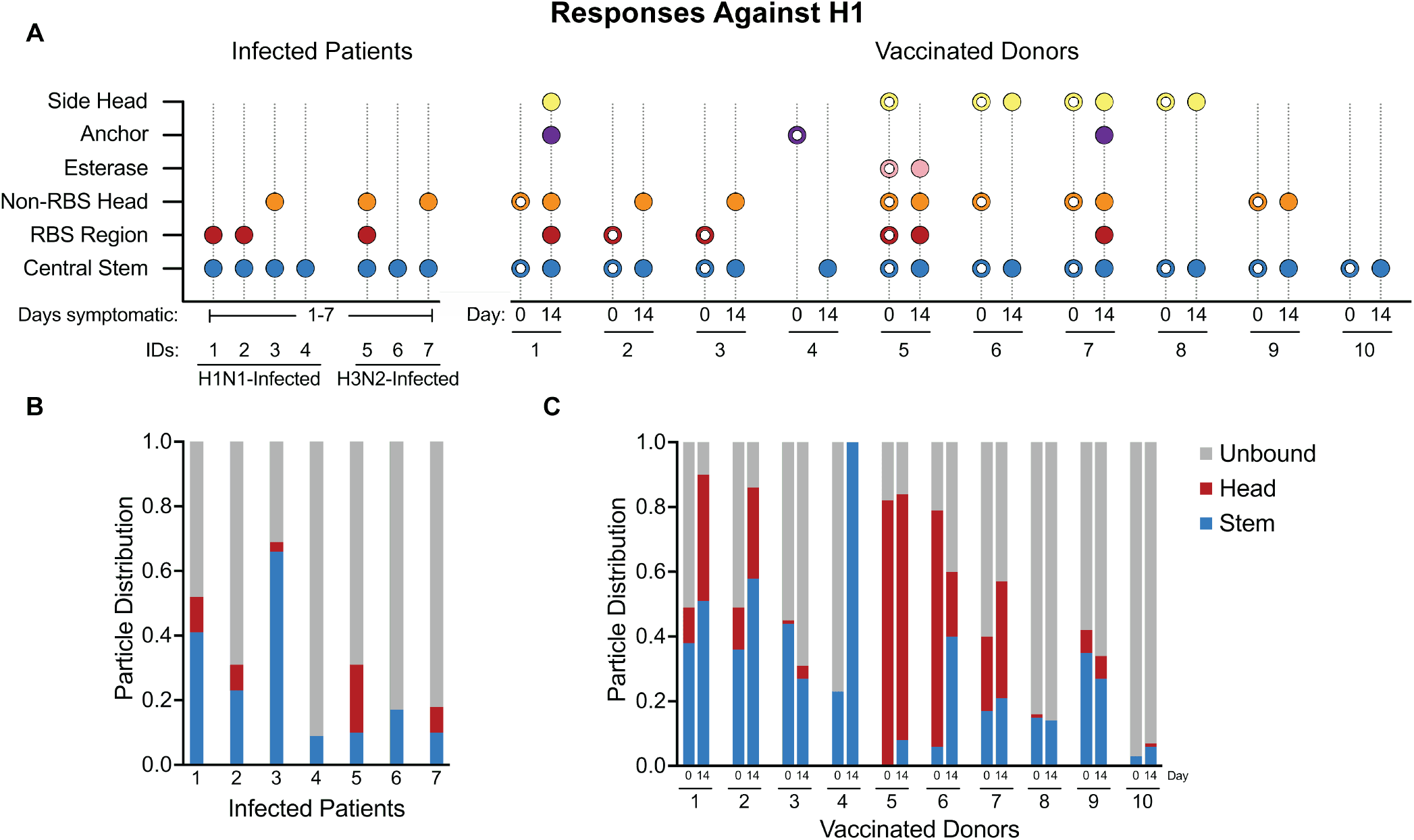
H1^+^ Fab responses reveal robust targeting of the central stem. **(A)** Visual summary of antibody responses against H1 present in vaccinated donors and infected patients. Open circles are used to differentiate vaccinated donor responses at Day 0 from responses observed at Day 14. **(B)** Stacked bar charts illustrating particle distribution in infected patient nsEMPEM datasets. **(C)** Stacked bar charts illustrating particle distribution in vaccinated donor nsEMPEM datasets.

**Figure 6:**
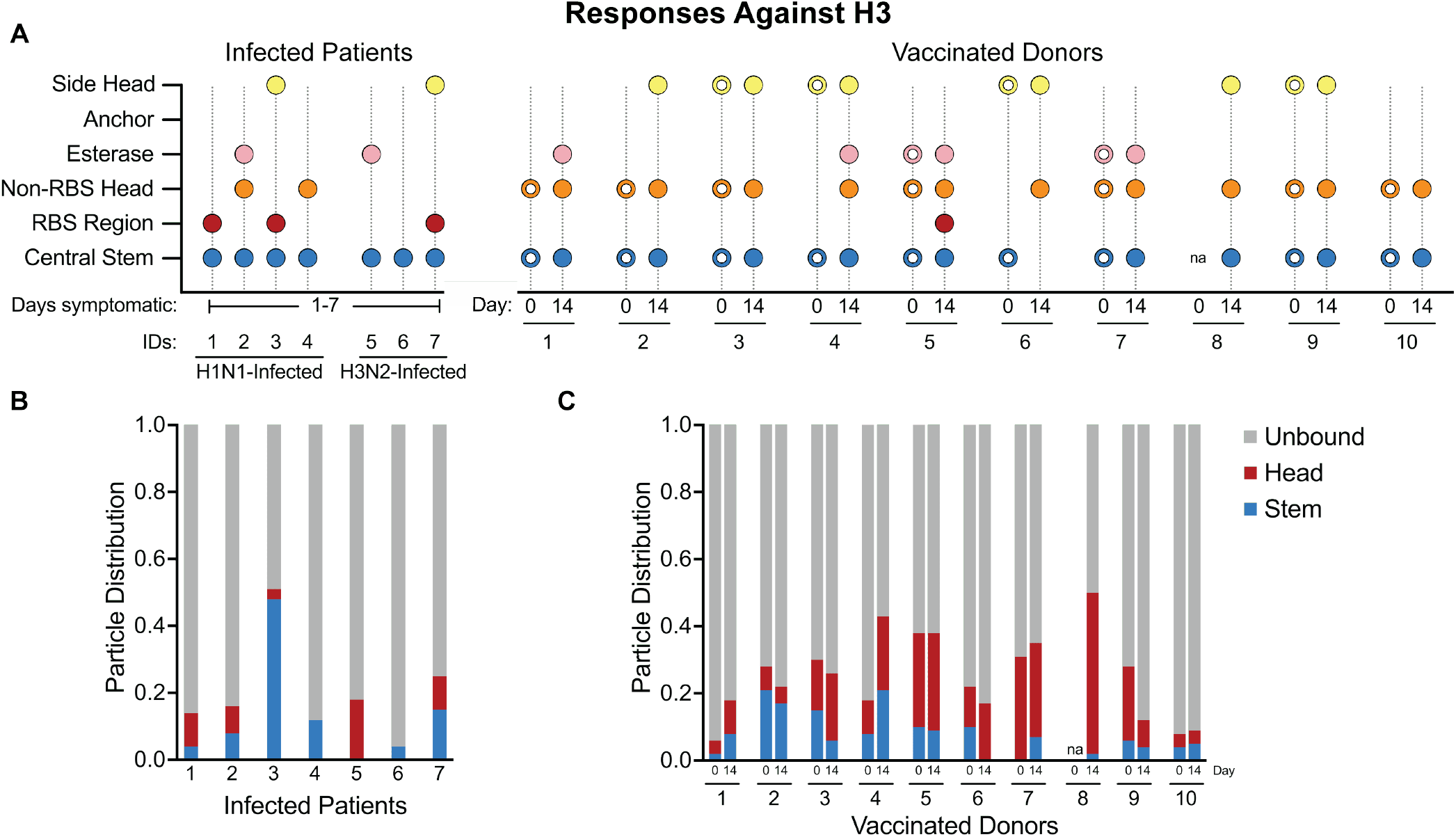
Central stem epitopes are a smaller fraction of particles observed in H3^+^ Fab responses. **(A)** Visual summary of antibody responses against H3 present in vaccinated donors and infected patients. Open circles are used to differentiate vaccinated donor responses at Day 0 from responses observed at Day 14. **(B)** Stacked bar charts illustrating particle distribution in infected patient nsEMPEM datasets. **(C)** Stacked bar charts illustrating particle distribution in vaccinated donor nsEMPEM datasets.

Although all vaccinated donors had consistent or greater HI titers on Day 14 compared to Day 0, only three had HI titers greater than the infected patients. In the context of ELISA results and particle distributions against H1, this suggests that while the infected patients may have had a lower number of HA^+^ antibodies or antibodies of lower affinity than those present in the vaccinated donors on Day 14, a greater proportion of these antibodies had HI activity—though they were present at levels insufficient for protection.

## Discussion

### Correlates of protection and central stem occupancy

Responses to the conserved central stem correlate with age, likely due to repeated exposure through vaccination and natural infection (38, 39). While numerous studies have demonstrated that immunization with engineered stem-targeting antigens can induce robust, broadly neutralizing antibody responses, not all stem-targeted responses successfully protect against acute infection in animal models (40). Our work supports this assessment by structurally mapping an abundance of stem-binding antibodies in plasma collected from emergency department patients presenting for care with symptomatic respiratory illness caused by IAV infection.

The apparent lack of protection conferred by the stem-binding antibodies observed in infected patients can be explained through one or more hypotheses: 1) the antibodies observed were protective but circulated at insufficient levels to prevent acute infection, 2) the antibodies targeted uncharacterized, non-protective sites on the stem epitope, 3) the antibodies targeted protective epitopes and circulated at high levels but were low affinity or incapable of engaging strong ADCC or ADCP and were therefore unable to effectively confer protection. While it is beyond the scope of this work, genetic variation in the immune system likely contributes to the outcomes observed.

Our ELISA data demonstrate a 4-fold difference in AUC between vaccinated patients at day 14 and infected patients sampled less than 14 days post-exposure, suggesting either a lower overall level of circulating anti-HA antibodies in the serum of the infected patients or drastically lower affinities (**Figure 4**). While there are many possible explanations for this observation, it is notable that the vaccinated subjects at Day 0 had AUCs similar to those of the infected patients.

EMPEM of infected patients showed dominance of stem responses. H1 stem responses were the predominant responses in six of the infected patients, and H3 stem responses were predominant in four of the infected patients (**Figures 5-6:B-C**). Interestingly, three of these patients were highly stem-responsive across both H1 and H3, raising the possibility of cross-reactive antibodies. However, nsEMPEM lacks the resolution to pursue this further.

EMPEM revealed that all vaccinated patients had prominent stem-targeting antibodies to both H1 and H3 at Day 0 and 14. Sixteen out of nineteen datasets also presented head-targeting antibodies on both days, demonstrating that LLPCs producing anti-head antibodies were also present before vaccination. nsEMPEM at Day 14 revealed epitopes unobserved at Day 0, which may indicate early GC-mediated or late extrafollicular plasmablast responses. Given the low resolution of nsEMPEM and lack of B-cell sequencing data, we are unable to deduce the contribution of GC-dependent somatic hypermutation on responses to epitopes present on Day 0 or Day 14. However, increases in the fraction of head- and stem-bound particles against H1 between Day 0 and Day 14 suggest that responses observed post-vaccination were either more prevalent or had higher affinity than those prevaccination—consistent with ELISA data (**Figures 5-6**).

While our data are insufficient to provide a complete, mechanistic explanation for the differences observed between patients infected with or vaccinated against IAV, they highlight interesting future research directions. The infected patients studied in our cohort appeared to have a bias towards the conserved HA stem, in accordance with previous work that found 84% of samples tested exhibited responses to the HA stem via ELISA (**Figures 2, 5-6**) (41). Notably, Patient 3 (age 79) had the highest proportion of stem-bound particles against H3 and the second highest to H1, reflecting a lifetime of repeated exposure to conserved central stem epitopes and/or the effects of early childhood imprinting.

Particle distributions and ELISAs suggest that these stem antibodies appear either less abundant or have lower affinity than those observed in vaccinated patients on Day 14.

### Epitope dynamics over time

Studies of immune responses in infected patients outside the context of challenge studies are complicated by the inability to determine temporal dynamics unambiguously. However, previous studies have demonstrated that influenza virus infection symptoms generally peak 2 to 3 days post-infection and that infection quickly leads to antigen-dependent B-cell activation, primarily in the respiratory tract draining lymph nodes (42– 44). Memory B cells with high affinity for influenza antigens undergo rapid clonal expansion and preferentially differentiate into extrafollicular short-lived plasmablasts, which dominate the early antibody response to primary infection and peak 5-7 days after symptom onset (43, 45, 46). Throughout this process, long-lived plasma cells (LLPCs) also secreted affinity-matured Abs against influenza antigen. Although cellular mechanisms are the primary contributors to viral clearance at this stage of infection, this blended Ab response contributes to clearance and regulates inflammatory responses associated with prolonged infection (46, 47).

As supported by previous work on this cohort of infected patients, antibody responses observed for all patients other than patient 6 are likely derived from LLPCs (45). Our results suggest that these responses failed to generate circulating antibody titers at protective levels, consistent with the patients’ decision to seek care in an emergency department. However, we found that infected patients had HAI titers equivalent to or better than those of the vaccinated donors. In the context of the comparatively low ELISA AUCs and the presence of RBS responses in nsEMPEM, the data suggest that infected patients did have circulating antibodies capable of robust HAI but not at levels sufficient to protect against symptomatic infection. This finding is consistent with our analysis of particle distributions, which indicate that head responses made up 10% or less of observed particles in all but one infected patient—Patient 5 had a more robust head response to both H1 and H3 (**Figures 5** and **6**).

The infected patients and vaccinated donors at Day 0 differed in the observable diversity of HA^+^ epitope responses. While the vast majority (10/14) of complexes analyzed for the infected cohort only presented top head and central stem antibodies, 13/20 complexes in the vaccinated cohort had at least one antibody targeting an epitope distinct from the top head and central stem.

Consistent with previous studies, we found that responses against H3 had lower titers than those against H1 (**Figures 4-6**) (48, 49). In vaccinated donors, ELISA AUCs against H1 were 46% higher than H3 at Day 0 and 35% higher at Day 14. Infected patients had AUCs against H1 58% higher than against H3. These trends are recapitulated in our analysis of particle distributions.

Unlike all other donors, Donor 6 lost a head response to H1 and a central stem response to H3 between Day 0 and Day 14. This finding may be a result of bias in the conditions for migration to a GC for SHM and affinity-maturation. Whereas high-affinity memory B-cells are preferentially recruited for extrafollicular responses, GC-recruitment is initiated by lower affinity interactions (50–52). Thus, the antibodies observed binding at Day 0 may not necessarily indicate the epitopes targeted by the memory B-cell cells that are most likely to be recruited to a GC.

### Limitations and future directions

Given low sample availability, we were unable to assess microneutralization titers for the infected patients, nor were we able to screen for Fc-effector function for infected patients or vaccinated donors. Moreover, while there is compelling evidence for uncharacterized stem epitopes, nsEMPEM requires complementary techniques to assess the specific identities, potential cross-reactivity, and the level of somatic hypermutation of the stem responses observed in this study. Additionally, since all nsEMPEM in this study was done using Fab, responses that depend on the avidity afforded by IgG have not been observed. Lastly, while purified IgG Fab is used for EMPEM, ELISAs, and HAI were conducted with plasma where IgA, IgM, and C1q may play a role in the trends observed. Some of these limitations could be overcome by conducting paired B-cell sequencing and high-resolution cryoEMPEM, which requires more Fab but can differentiate distinct antibodies targeting overlapping epitopes, as previously reported for an engineered HIV envelope glycoprotein (36). As sample availability for EMPEM studies increases and methodology improves, we aim to increase our capacity to incorporate these complementary approaches in future work.

## Conclusion

Our nsEMPEM analysis of infected patients and vaccinated donors found that central stem responses accounted for the majority of particles observed in H1 data sets. H3 responses were more evenly distributed and contained a greater fraction of unbound HA particles compared to H1 data sets. While it is difficult to ascertain the full diversity of responses to HA at the negative stain level—what appear to be single responses in negative stain can be an average of multiple responses with overlapping footprints observable at higher resolution— responses to H3 generally targeted a greater number of epitopes than responses to H1, despite lower ELISA AUCs and a lower proportion of bound particles. In the context of previous EMPEM and B-cell sequencing studies, this work continues to highlight the range of HA^+^ responses present in circulating IgG repertoires from person to person and emphasizes the importance of expanding sample size in future work (19, 35, 53, 53–56). While additional functional studies are necessary to explore this observation, the abundance of central stem responses among emergency department patients emphasizes the need to carefully tailor rational vaccine designs to elicit protective responses to this highly conserved epitope. As our capacity to conduct structural and single-cell analyses grows, we aim to interrogate differences between protective and non-protective responses to influenza infection at scales akin to those of systems serology.

## Acknowledgements

The Ward lab was funded by an award provided by the Human Vaccines Project, Inc. The Crowe lab was funded by an award provided by the Human Vaccines Project, Inc, HHS Contract 75N93019C00074, and NIAID grant U01 AI150739. The Mudd laboratory was supported by a grant from the NIH R01AI173203 and by a grant from the Emergency Medicine Foundation. The Vanderbilt Vaccine Research Program received funding from the Human Vaccine Project. Rachael M. Wolters is funded by K01 OD036063. We thank Hailee Perrett for assistance producing Figure 1 and acknowledge the collective effort of the KIRV and EDFLU clinical study teams for collecting, storing, and distributing sera and plasma samples used in this work. We also acknowledge the patients and volunteers who chose to make their biospecimens available to the study teams at point-of-care (EDFLU) or through study enrollment (KIRV).

## Author contributions

ANL and AJR produced recombinant HA; ANL, AJR, STR, JH, ATP, and AMJ conducted EMPEM experiments; ANL and RMW performed serology; RMW, KW, and JEC provided recombinant influenza virus; PAM collected infected patient samples; CBC and SY enrolled study participants and processed trial samples; ANL wrote the manuscript; JH, PAM, JEC, and ABW conceived experimental questions; all authors edited and reviewed the manuscript prior to submission.

## Competing interest statement

J.E.C. has served as a consultant for Luna Labs USA, Merck Sharp & Dohme Corporation, Emergent Biosolutions, a former member of the Scientific Advisory Boards of Gigagen (Grifols), of Meissa Vaccines, and BTG International, is founder of IDBiologics and receives royalties from UpToDate. The laboratory of J.E.C. received unrelated sponsored research agreements from AstraZeneca, Takeda Vaccines, and IDBiologics during the conduct of the study. CBC has served as a consultant to Sanofi Pasteur, Pfizer, Moderna, GSK, TDCowen, Debiopharm, and CommenseBio and receives royalties from UoToDate. All other authors declare they have no competing interests.

## Materials and Methods

### Vaccinated donor sample collection

Blood samples were obtained from healthy human donors who had received the quadrivalent cell-based influenza vaccine, with all procedures approved by the Vanderbilt University Medical Center Institutional Review Board. Peripheral blood was collected using standard venipuncture techniques into either EDTA-coated tubes for plasma or serum-separator tubes for serum. Post-centrifugation, plasma and serum were aliquoted into cryovials, avoiding contamination with cellular components, and stored at -80°C until analysis. Each sample was labeled with a unique identifier for traceability, and storage conditions were continuously monitored to maintain sample integrity.

### Infected patient sample collection

Infected subjects were enrolled into the EDFLU study (45). We approached individuals over the age of 18 with a positive clinical influenza real-time reverse-transcription polymerase chain reaction test (Cepheid Xpert Flu/RSV) who presented for care at the Emergency Department of Barnes Jewish Hospital (Saint Louis, Missouri, USA). Enrolled participants must have experienced influenza symptoms within the 24 hours prior to enrollment. We obtained written informed consent from the subjects or their legally authorized representative. The EDFLU study was approved by the Washington University in Saint Louis Institutional Review Board (approval numbers 2017-10-220 and 2018-08-115). Blood was collected into EDTA anticoagulated tubes and plasma fractions were obtained by centrifugation within 8 hours of phlebotomy. Plasma was stored at -80°C or colder until analysis.

### Protein expression and purification

Recombinant H1 from A/Michigan/45/2015 and H3 from A/Singapore/INFIMH-009/2016 and A/North Carolina/04/2016 were expressed and purified from HEK293F (Thermo Fisher) cells. Cells were transfected at a ∼1.0 x 10^6^ cells/mL using PEIMAX with 0.5 to 0.75 mg/L of DNA at a ratio of 3:1 PEI:DNA. Cells were cultured at 37°C, 8% CO_2_, and shaken at 125 rpm in 293FreeStyle expression medium (Life Technologies). 5-6 days after transfection, cells were pelleted at 8,000 x *g* for 20 min and the supernatant was filtered through 0.22-um filter. Nickel nitrilotriacetic acid (Qiagen) agarose beads were washed in TBS and added to filtered supernatant at a ratio of 2-3 mL resin/L and incubated on a rotator over-night at 4°C. Samples were subsequently run over a gravity flow column (Qiagen) and washed with TBS and 20 mM imidazole pH 8.0. Proteins were eluted with 250 mM imidazole and buffer exchanged into TBS three times on a 30 kDa concentrator (Amicon). Affinity-purified HA was purified using size exclusion chromatography over a Superdex 200 Increase 10/300 column (GE Healthcare). Fractions corresponding to trimeric HA were pooled, concentrated, and buffer exchanged to TBS using 30 kDa Amicon concentrators.

### IgG purification, digestion, and complexing

Serum or plasma samples were heat-inactivated by incubating 1 mL aliquots in a 55°C water bath for 1 h. HI samples were incubated with 1 mL of IgG Fc Capture Select (Thermo Fisher) resin at 4C for 20 to 48 h to bind IgG. Resin was pelleted by centrifuging in an aerosol-tight centrifuge for 5 m at 200 x *g*. IgG-depleted samples were decanted and resin was washed three times with PBS. Papain was activated by incubating freshly prepared Tris-EDTA, L-cysteine, and papain at a final concentration of 100 mM Tris, 2 mM EDTA, 10 mM L-cysteine, and papain 1 mg/mL at 37°C for 15 min. Bound IgG were digested on-resin to elute Fab by incubating at 37°C for 4-5 h with 400 μL of activated papain. Digestion was halted by adding iodoacetamide to a final concentration of 0.03 M. Fab was buffer exchanged on a 10 kDa Amicon concentrator and purified through size exclusion chromatography with a Superdex 200 increase 10/300 column (GE Healthcare). Complexes were generated by incubating Fab and antigen overnight at a 50:1 ratio and purified through size exclusion chromatography with a Superdex 200 increase 10/300 column (GE Healthcare).

### Negative stain EMPEM

To prepare nsEMPEM grids, 3 μL of purified immune complexes were applied to glow-discharged, carbon-coated 400 mesh copper grids in at a concentration of ∼20 μg/mL (Electron Microscopy Services). Excess sample was blotted with Whatman filter paper and 3 μLs of 2% w/v uranyl formate were immediately applied to stain complexes for 60 s twice. Samples were imaged on either a Tecnai T20 (FEI) with an Eagle CCD 4k camera (FEI) at 200 kV, 62,000 × magnification, and 1.77 Å/pixel; a Talos 200C with a Falcon II direct electron detector and a CETA 4k camera (FEI) at 200 kV, 73,000 × magnification, and 1.98 Å/pixel; or a Tecnai Spirit T12 (FEI) with a CMOS 4k camera (TVIPS) at 120 kV, 52,000 × magnification, and 2.06 Å/pixel. Micrographs were collected using Leginon. For each complex, 100k to 400k particles were picked and stacked using Appion and subsequently processed to generate 2D classes and 3D reconstructions using Relion 3.0 (57–60). UCSF Chimera was used to generate composite 3D reconstructions (61). Complexes that could not be resolved in 3D were presented as false-colored 2D classes. For 3D reconstructions where an epitope was clearly targeted, but the Fab region was poorly resolved, our previous work, 2D classes, and the partial Fab density was used to assign a cartoon Fab to the targeted region as a best possible prediction as in (35).

### Analyzing EM particle distribution

Following 2D classification with Relion 3.0, unbound HA side views, head-bound views, and stem-bound views were counted and logged. Particle distribution charts were generated using GraphPad Prism.

### Plasma ELISAs

High-binding 96-well plates (Perkin Elmer SpectraPlate 96 HB, #6005609) were coated overnight at 4°C with HA proteins at a concentration of 2 μg/mL in PBS (50 μL/well). Plates were washed three times using PBS containing 0.1% Tween 20 (PBS-T). Wells were blocked using 200 μL of PBS-T plus 5% bovine serum albumin (BSA) (insert) overnight at 4°C or 2 h at room temperature. Plates were then washed three times with PBS-T. Plasma was heat inactivated at 56°C for 1 h. 2-fold serial dilutions were prepared in PBS-T plus 1% BSA in non-binding 96-well plates (Greiner Bio-One, #655901) in triplicate at a starting dilution of 1:50 and a final volume of 60 μL/well. 50 μLs of dilutions and controls were transferred to antigen-coated plates and incubated for 1 h at room temperature. Wells were washed three times with 200 μL of PBS-T. Peroxidase-conjugated goat secondary anti-human IgG F(ab′)_2_ fragment specific (Jackson Laboratories, 109-035-097) antibody was diluted 1:5000 in PBS-T plus 1% BSA and 100 μL of secondary antibody solution was added to all wells and incubated for 1 h at room temperature. Plates were washed three times with 200 μL of PBS-T and developed using 100 μL/well of o-phenylenediamine dihydrochloride (OPD, Sigma P8287) for 10 min. The reaction was stopped with 100 μL of 1 N sulfuric acid and then read at an absorbance of 450 nm with a Synergy H1 hybrid multimode microplate reader (BioTek). Area under the curve analysis was conducted using GraphPad Prism.

### HA inhibition assay

HAI assays to determine IC100 values. For the HAI assay, we used turkey red blood cells (RBCs) in Alsever’s solution (LAMPIRE Biological Products, #7209403) diluted to 1% in D-PBS (Corning, #21031CM). 50 μL of 4 hemagglutination units of virus were incubated with 50 μL of 3-fold dilutions of the serum in D-PBS, and 50 μL of RBCs for 1 hr at room temperature. The HAI titer was defined as the highest dilution of antibody that inhibited hemagglutination of red blood cells. Each dilution was performed with technical duplicates and in biological duplicate.

### Data availability

EM maps for all complexes have been deposited in the electron microscopy data bank (EMDB) and all accession codes have been listed in supplemental material (**Table S1**).

## Supplementary Material

**Figure S1:**
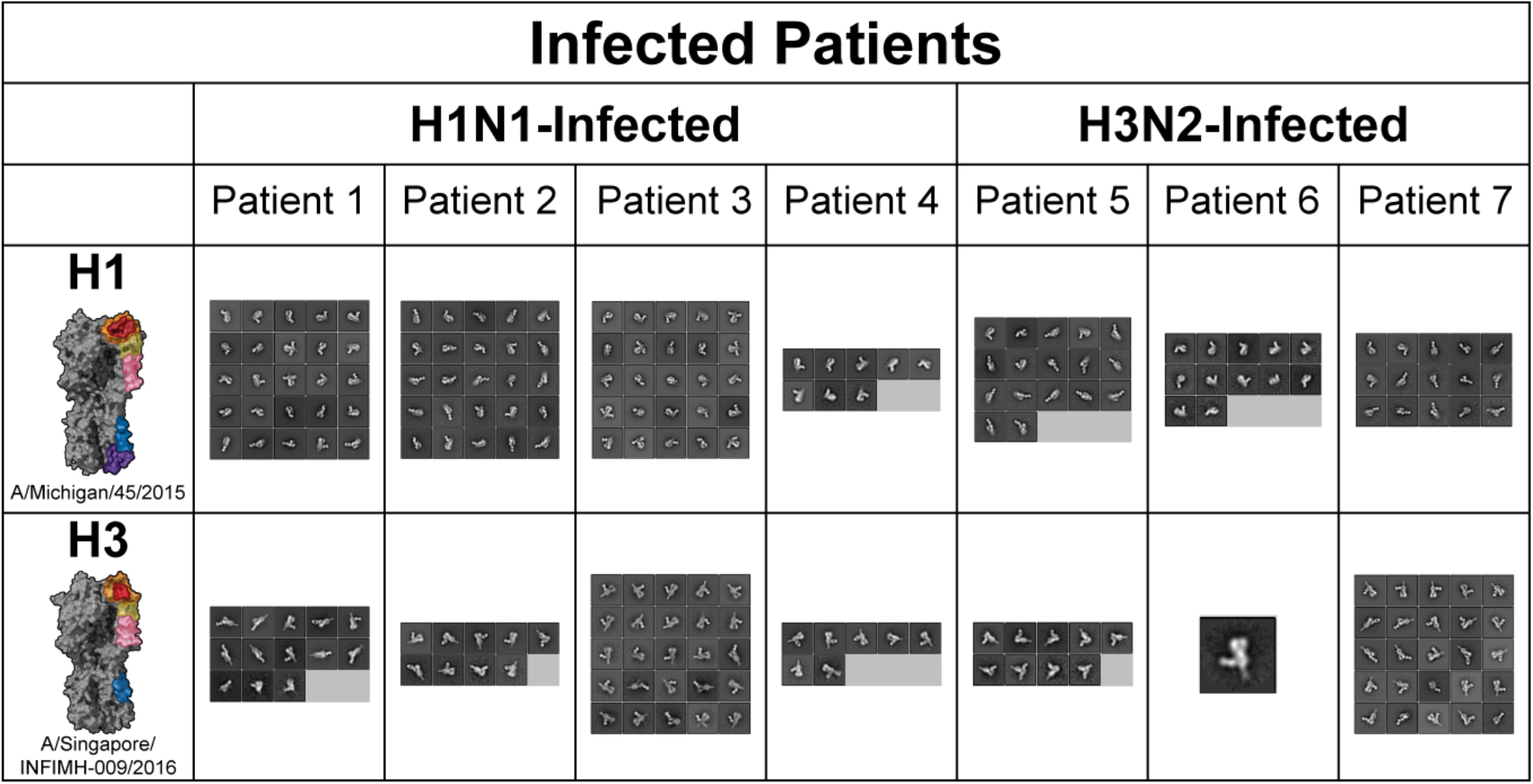
Representative 2D classes of infected patient Fabs bound to H1 and H3. Classes are representative of particles incorporated in final 3D reconstructions and/or cartoon Fab representations.

**Figure S2:**
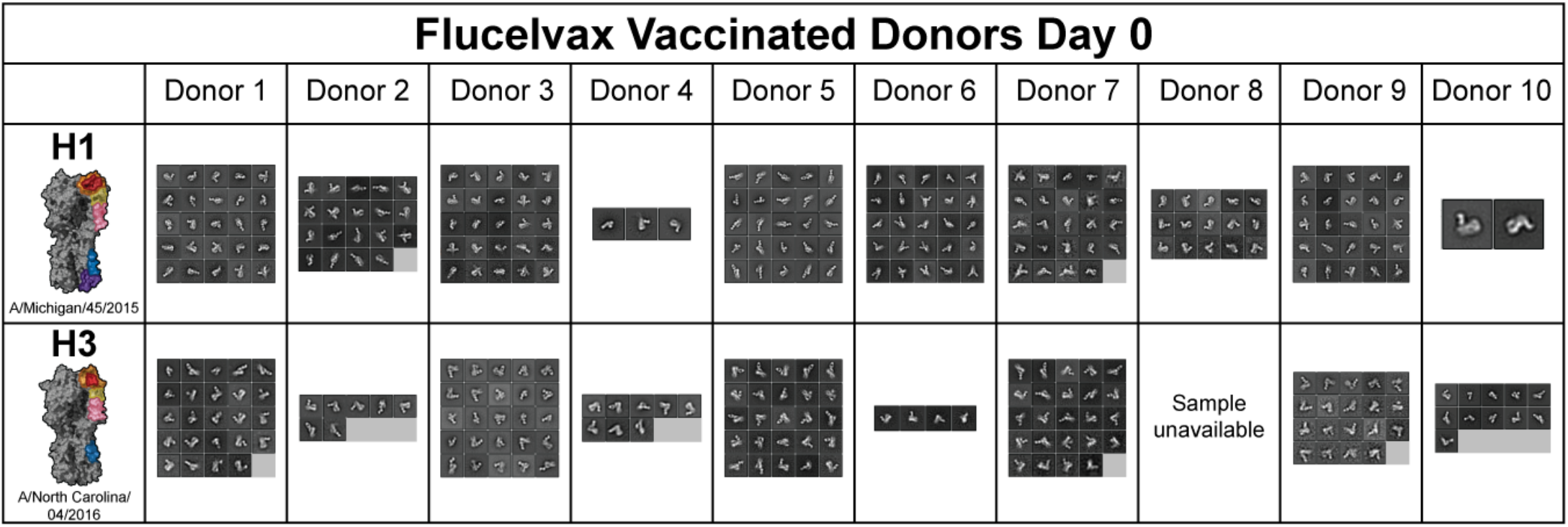
Representative 2D classes of vaccinated donor Fabs bound to H1 and H3 at Day 0. Classes are representative of particles incorporated in final 3D reconstructions and/or cartoon Fab representations.

**Figure S3:**
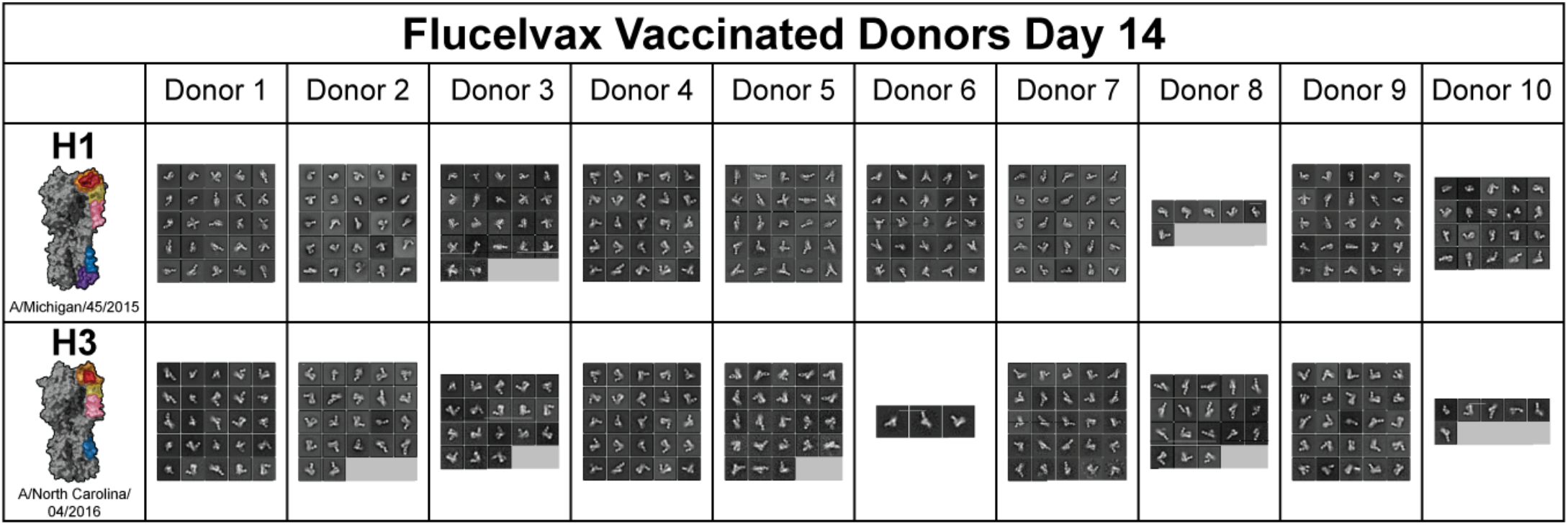
Representative 2D classes of vaccinated donor Fabs bound to H1 and H3 at Day 14. Classes are representative of particles incorporated in final 3D reconstructions and/or cartoon Fab representations.

**Table S1:**
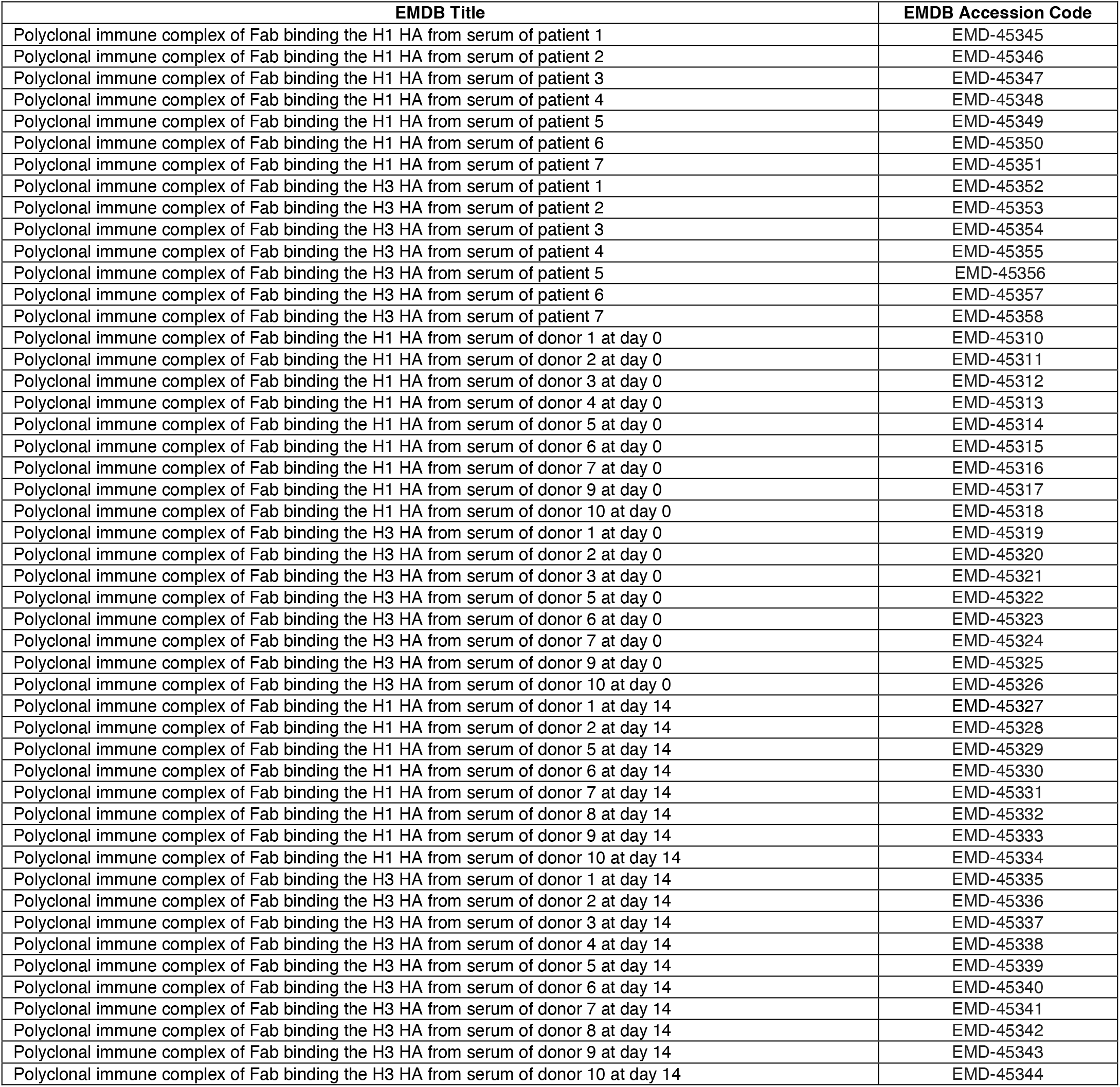
EMDB deposition ID. Global refinements, half maps, and 3D classes used to generate all negative stain EM reconstructions presented in this manuscript have been deposited to the EMDB and can be accessed using the data provided in the table below.

